# New roles for an old cyclin in control of cell cycle entry and cell size

**DOI:** 10.1101/2022.08.12.503734

**Authors:** Amanda Brambila, Beth E. Prichard, Jerry T. DeWitt, Douglas R. Kellogg

## Abstract

Entry into the cell cycle in late G1 phase occurs only when sufficient growth has occurred. In budding yeast, a cyclin called Cln3 is thought to link cell cycle entry to cell growth. Cln3 accumulates during growth in early G1 phase and eventually triggers accumulation of late G1 phase cyclins that drive cell cycle entry. All current models for cell cycle entry assume that expression of late G1 phase cyclins is initiated at the transcriptional level. Current models also assume that the sole function of Cln3 in cell cycle entry is to promote transcription of late G1 phase cyclins, and that Cln3 works solely in G1 phase. Here, we show that cell cycle-dependent expression of late G1 phase cyclins does not require cell cycle-dependent transcription. Moreover, Cln3 can influence accumulation of late G1 phase cyclin proteins via post-transcriptional mechanisms. Finally, we show that Cln3 has functions in mitosis that strongly influence cell size. Together, these discoveries reveal surprising new functions for Cln3 that challenge current models for control of cell cycle entry and cell size.

## Introduction

The decision to enter a new round of cell division is amongst the most consequential decisions in the life of a cell. Entry into the cell cycle occurs only when sufficient growth has occurred and only when there are sufficient nutrients for further growth. In animal cells, cell cycle entry is also controlled by growth factors, which ensure that cell division occurs at an appropriate time and place. Defects in signals that control cell cycle entry are a primary cause of cancer.

The mechanisms that initiate cell cycle entry are poorly understood (Rubin *et al*., 2020). In budding yeast, decades of work led to a canonical model in which a cyclin called Cln3 appears in early G1 phase and activates Cdk1 (Carter and Sudbery, 1980; Cross, 1988, 1989; Nash *et al*., 1988; Hadwiger *et al*., 1989; Richardson *et al*., 1989; Tyers *et al*., 1992, 1993). The Cln3/Cdk1 complex then directly phosphorylates and inactivates Whi5, a transcriptional repressor that binds and inhibits two transcription factors, referred to as SBF and MBF, that drive transcription of late G1 cyclins, as well as hundreds of additional genes (Costanzo *et al*., 2004; de Bruin *et al*., 2004; Iyer *et al*., 2001; Koch *et al*., 1996; Nasmyth and Dirick, 1991; Ferrezuelo *et al*., 2010; Tyers *et al*., 1993; Wijnen *et al*., 2002; Jorgensen *et al*., 2002; Spellman *et al*., 1998; Bean *et al*., 2005). Thus, Cln3-dependent inactivation of Whi5 has been thought to initiate transcription of late G1 cyclins, which is the critical molecular event that marks cell cycle entry. Genetic analysis suggests that Cln3 and Whi5 link cell cycle entry to cell growth. For example, loss of *WHI5* causes premature cell cycle entry before sufficient growth has occurred, leading to a reduced cell size (Jorgensen *et al*., 2002; Costanzo *et al*., 2004; de Bruin *et al*., 2004). Similarly, loss of Cln3 causes delayed cell cycle entry and increased cell size, while overexpression of Cln3 leads to premature cell cycle entry and reduced cell size (Sudbery *et al*., 1980; Cross, 1988, 1989; Nash *et al*., 1988; Tyers *et al*., 1993; Jorgensen and Tyers, 2004).

The canonical model has provided an important framework for analysis of cell cycle entry and has strongly influenced models for how cell growth influences cell cycle entry. However, a number of observations cannot be explained by the model. For example, over-expression of Cln3 causes a large reduction in the size of *whi5*Δ cells, which indicates that Cln3 has critical targets other than Whi5 (Costanzo *et al*., 2004; Wang *et al*., 2009). Furthermore, recent work has shown that Cln3 is not required for phosphorylation of Whi5 in vivo (Bhaduri *et al*., 2015; Kõivomägi *et al*., 2021). Rather, a recent study suggested a revised model in which Cln3/Cdk1 binds to SBF promoters and directly phosphorylates RNA polymerase to stimulate transcription (Kõivomägi *et al*., 2021). This revised model, as well as the canonical model, presume that Cln3 functions entirely at the transcriptional level to promote synthesis of late G1 cyclin proteins. However, this has never been directly tested.

A further concern regarding current models is that they assume that Cln3 exerts all of its key functions in G1 phase, especially with respect to control of cell size. However, there are two peaks of Cln3 protein during the cell cycle – one in G1 phase and a second in mitosis (Landry *et al*., 2012; Zapata *et al*., 2014; Litsios *et al*., 2019). The functions of Cln3 during mitosis are unknown. Since most growth of a yeast cell occurs during bud growth in mitosis (Leitao and Kellogg, 2017), it is possible that mitotic functions of Cln3 play a major role in Cln3’s ability to influence cell size. No previous experiments have tested for mitotic functions of Cln3.

Here, we tested key aspects of current models for cell cycle entry. We also tested whether Cln3 has functions in mitosis. Together, the data indicate that Cln3 can strongly influence accumulation of late G1 cyclins via post-transcriptional mechanisms, and that Cln3 executes functions during mitosis that influence cell size. Together, these observations significantly expand the range of possible models for cell size control and the functions of Cln3.

## Results

### Cln3 can influence production of Cln2 protein via Whi5-independent mechanisms

Previous studies examined how Cln3 and Whi5 influence Cln2 mRNA levels. Here, we examine how Cln3 and Whi5 influence production of Cln2 protein, which is the critical output of the mechanisms that drive cell cycle entry.

We first set out to test how Cln3 and Whi5 contribute to regulation of Cln2 protein expression. Recent work suggests that Cln3/Cdk1 does not phosphorylate Whi5 (Bhaduri *et al*., 2015; Kõivomägi *et al*., 2021); however, it remains possible that Cln3 drives dissociation of Whi5 from SBF via other mechanisms that could make a substantial contribution to expression of Cln2 protein. Therefore, we tested whether Cln3 can influence production of Cln2 protein via mechanisms that are independent of Whi5. To do this, we analyzed the effects of loss- or gain-of-function of Cln3 on production of Cln2 protein in *whi5*Δ cells. If Cln3 influences expression of Cln2 protein primarily via Whi5-dependent mechanisms, then loss- or gain-of-function of Cln3 should have little effect on Cln2 protein expression in *whi5*Δ cells. In these experiments and additional experiments that follow we detected Cln2 using multiple N-terminal and C-terminal tags to ensure that results are not influenced by the location or composition of the tag. All tagged versions of Cln2 gave similar results, although detection of phosphorylated forms of Cln2 that have reduced electrophoretic mobility differed between the tags.

We used *cln3*Δ to test the effects of loss of Cln3 function. To achieve gain of Cln3 function, we utilized the *cln3*-Δ*177* allele, which lacks C-terminal PEST sequences that target it for rapid turnover via ubiquitin-dependent proteolysis (Nash *et al*., 1988). Previous work found that *cln3*-Δ*177* results in a 10-fold increase in Cln3 protein levels, as well as a large decrease in cell size (Sudbery *et al*., 1980; Tyers *et al*., 1992). We synchronized wild type, *whi5*Δ, *whi5*Δ *cln3*-Δ*177* and *whi5*Δ *cln3*Δ cells in G1 phase using mating pheromone and assayed production of *Cln2-3XHA* during the cell cycle at 10 minute intervals (Figures 1A,B). Loss of *WHI5* accelerated production of Cln2 protein by approximately 10 minutes relative to wild type, as expected. Overexpression of Cln3 accelerated production of Cln2 protein in *whi5*Δ cells, while *cln3*Δ delayed production of Cln2 in the *whi5*Δ cells. Thus, Cln3 can strongly influence production of Cln2 protein via mechanisms that are independent of Whi5. Furthermore, Cln2 protein expression remained strongly periodic in *whi5*Δ *cln3*Δ cells, which indicates that Cln3 is not required for robust cell cycle-dependent expression of Cln2 protein.

**Figure 1:**
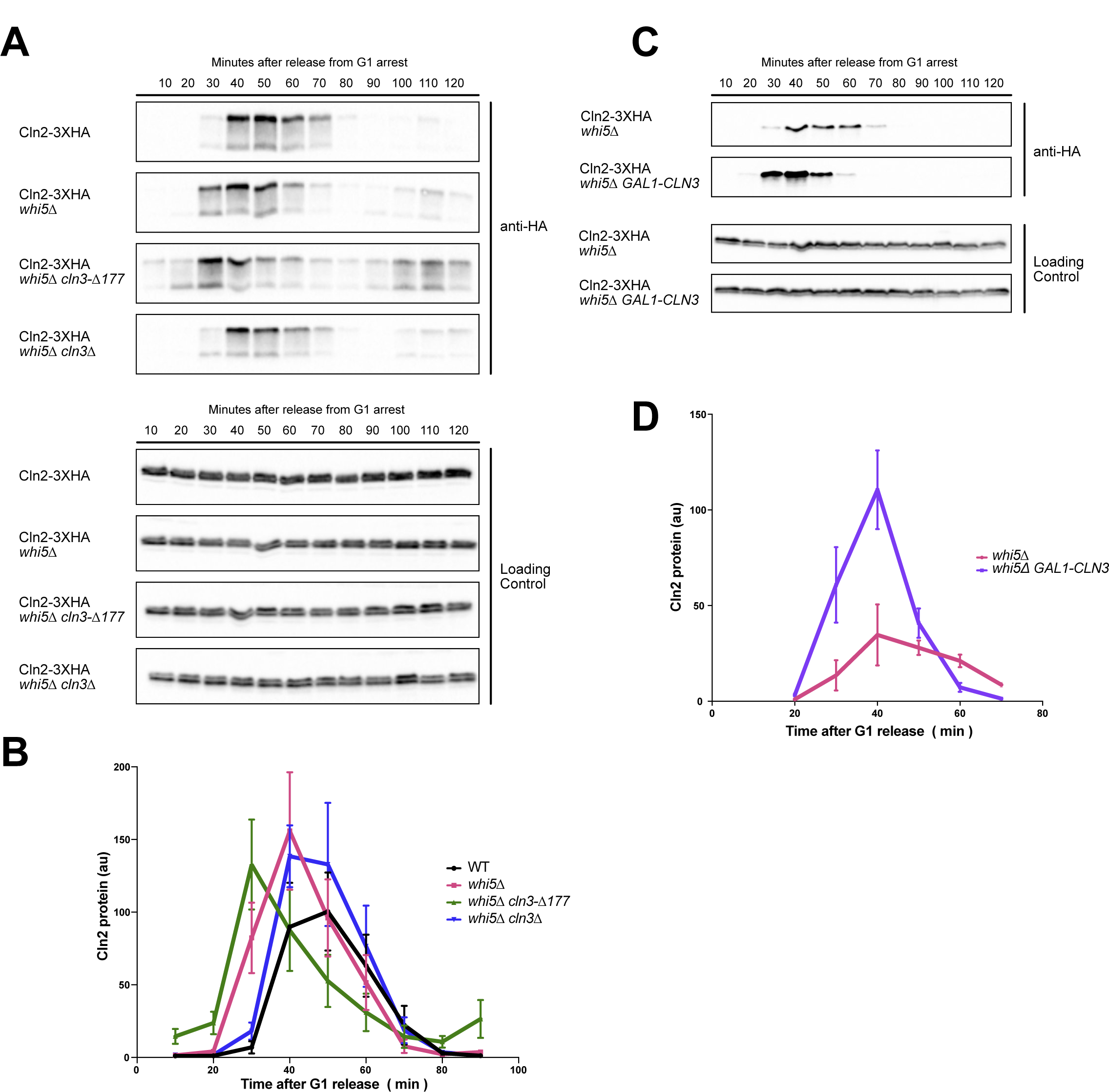
Cln3 can influence production of Cln2 protein via Whi5-independent mechanisms. **(A)** Cells were grown in YPD medium overnight and synchronized in G1 phase using alpha factor. Cells were released from G1 arrest and samples were collected every 10 minutes to assay for Cln2-3XHA protein levels by western blotting using 12CA5 mouse monoclonal anti-HA. An anti-Nap1 antibody was used as the loading control. **(B)** Quantification of data in panel A. Cln2 protein levels were normalized against loading control. Error bars represent the standard error of the mean for 3 biological replicates. **(C)** Cells were grown in 2% YPGE medium overnight and synchronized in G1 phase using mating pheromone. 2% galactose was added to cultures at 40 minutes prior to release from G1 arrest. Cells were released from G1 arrest into YP medium containing 2% galactose and samples were collected every 10 minutes to assay for Cln2 protein levels by western blotting using 12CA5 mouse monoclonal anti-HA. **(D)** Quantification of data in panel C. Error bars represent standard error of the mean for 3 biological replicates.

A previous study concluded that Cln3/Cdk1 promotes transcription of *CLN2* via direct phosphorylation of RNA polymerase (Kõivomägi *et al*., 2021). The most straightforward interpretation of this model would predict that loss of Cln3 in *whi5*Δ cells should cause reduced transcription of *CLN2* and a corresponding reduction in Cln2 protein levels. However, we found that *cln3*Δ did not cause a reduction in peak Cln2 protein levels in *whi5*Δ cells (Figure 1B, compare *whi5*Δ and *whi5*Δ *cln3*Δ). Furthermore, *cln3*Δ *whi5*Δ cells express slightly more Cln2 protein than wildtype cells and the timing of expression of Cln2 protein in *whi5*Δ *cln3*Δ cells is indistinguishable from wild type cells (Figure 1B).

A concern with the use of *cln3*Δ and *cln3*-Δ*177* is that they cause substantial cell size defects that could indirectly influence the timing of cell cycle entry. Therefore, as a further means of testing the effects of Cln3 overexpression, we utilized cells that express an extra copy of wild type *CLN3* from the inducible *GAL1* promoter, which allowed us to test the immediate effects of increased Cln3 levels in otherwise normal cells. Wild type and *GAL1-CLN3* cells were arrested in G1 phase in media containing 2% glycerol and 2% ethanol to repress transcription of *GAL1-CLN3*. Expression of *GAL1-CLN3* was induced with 2% galactose 30 minutes before release from the G1 phase arrest. We again found that overexpression of Cln3 accelerated production of Cln2 protein in *whi5*Δ cells (Figure 1C,D). The effects of full length *CLN3* expressed from the *GAL1* promoter were stronger than the effects of *cln3*-Δ*177*, as *GAL1-CLN3* caused a larger increase in the amount of Cln2 protein (compare Figures 1B and 1D). This may be due to the fact that *cln3*-Δ*177* lacks a nuclear localization sequence (Edgington and Futcher, 2001; Miller and Cross, 2001).

Together, these data indicate that Cln3 can strongly influence Cln2 protein expression via Whi5-independent mechanisms, and that robust periodic expression of Cln2 does not require Cln3 or Whi5. The results also indicate that Cln3-dependent phosphorylation of RNA polymerase is unlikely to make a major contribution to the mechanisms responsible for cell cycle-dependent expression of Cln2 protein.

### Cln3 can influence levels of Cln2 protein via post-transcriptional mechanisms

Cln3 could control transcription of *CLN2* via mechanisms that are independent of Whi5, but dependent upon other features of the *CLN2* promoter. Alternatively, Cln3 could influence production of Cln2 protein via post-transcriptional mechanisms, which has never been tested. To distinguish these possibilities, we created strains in which transcription of *CLN2* is controlled by the *MET25* promoter, thereby eliminating normal control of *CLN2* transcription. Previous studies have shown that SBF and MBF cannot be detected at the *MET25* promoter and that overexpression of Cln3 does not influence transcription of *MET25* (Iyer *et al*., 2001; Ferrezuelo *et al*., 2010). The *MET25* promoter drives a basal level of transcription when methionine is present in the growth medium, which we used to express *CLN2*.

We found that *cln3*-Δ*177* advanced expression of Cln2 protein in *MET25pr-CLN2* cells, whereas *cln3*Δ caused delayed and reduced expression of Cln2 protein. (Figures 2A,B). We carried out a further test using a different heterologous promoter. In this case, we used the *YPK1* promoter, which does not undergo cell cycle-dependent transcription and does not bind SBF or MBF. In addition, the Ypk1 protein is expressed at constant levels throughout the cell cycle and across different carbon sources (Alcaide-Gavilán *et al*., 2018; Lucena *et al*., 2018). We found that *GAL1-CLN3* accelerated production of Cln2 protein expressed from the *YPK1* promoter (Figures 2C,D). As in Figure 1, the effects of *GAL1-CLN3* appeared to be stronger than the effects of *cln3*-Δ*177*.

**Figure 2:**
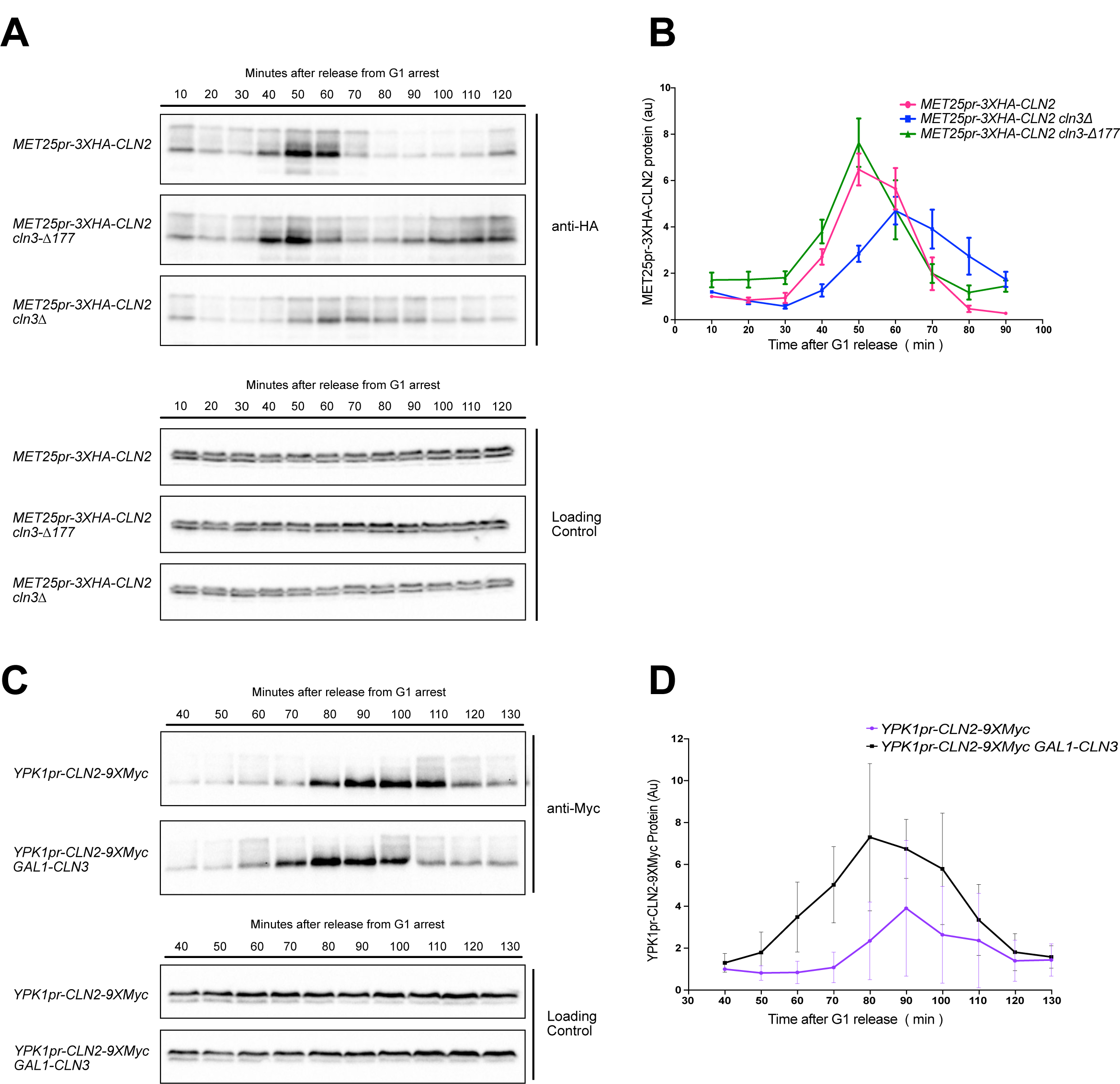
Cln3 can influence levels of Cln2 protein via post-transcriptional mechanisms. (A) Cells were grown in YPD medium overnight and synchronized in G1 phase using alpha factor. Cells were released from G1 arrest and samples were collected every 10 minutes to assay for MET25pr-3XHA-Cln2 protein levels by western blotting using 12CA5 mouse monoclonal anti-HA. An anti-Nap1 antibody was used as the loading control. (B) Quantification of the data in panel A. Plots were generated by normalizing MET25pr-3XHA-CLN2 to loading control. Error bars represent the standard error of the mean for 4 biological replicates. (C) Cells were grown in YPGE medium overnight and synchronized in G1 phase using alpha factor. 2% galactose was added to cultures 45 minutes prior to release from G1 arrest. Cells were then released from G1 arrest into YP medium containing 2% galactose. Samples were collected every 10 minutes, from 40 minutes after release from G1 phase to 130 min after release from G1 phase. Western blotting was used to assay protein levels for Cln2 using anti-Myc and anti-Nap1 for loading control. (D) Quantification of the data in panel D. Plots were generated by normalizing pYPK1-Cln2-9Myc to loading control. Error bars represent the standard error of the mean for 3 biological replicates.

These results show that Cln3 can influence expression of Cln2 protein via mechanisms that are completely independent of the *CLN2* promoter. Furthermore, Cln2 protein expressed from heterologous promoters showed strong cell cycle periodicity, which provides further evidence that post-transcriptional mechanisms play a substantial role in the mechanisms that drive periodic expression of Cln2 protein. These results also provide further evidence that Cln3-dependent phosphorylation of RNA polymerase at *CLN2* promoters is unlikely to play a major role in controlling periodic expression of Cln2 protein. Previous studies have shown that constitutive expression of *CLN1* from the very strong *GAL1* promoter can rescue the inviability of *cln1*Δ *cln2*Δ *cln3*Δ cells (Richardson *et al*., 1989). Although *GAL1-CLN1 cln2*Δ *cln3*Δ cells have an abnormal morphology, this observation shows that the essential functions of late G1 phase cyclins are not dependent upon regulation of their transcription, which further suggests that post-transcriptional mechanisms play an important role in regulating the activity of late G1 phase cyclins.

### Cln3 can not influence Cln2 protein levels in *swi6*Δ cells

Cln3 is thought to influence transcription of late G1 cyclins via two transcription factors, known as SBF and MBF. Both are dimers that include a DNA binding subunit and a shared subunit called Swi6 (Andrews and Herskowitz, 1989; Andrews and Moore, 1992; Siegmund and Nasmyth, 1996). SBF and MBF bind the promoters of late G1 phase cyclins, as well as hundreds of additional genes. Loss of Swi6 causes a large reduction in transcription of *CLN2*, as well as delayed cell cycle entry and a large increase in cell size (Nasmyth and Dirick, 1991; Wijnen *et al*., 2002). Key phenotypic effects of *CLN3* over-expression are not seen in *swi6*Δ cells. For example, overexpression of *CLN3* does not cause reduced cell size in *swi6*Δ cells and fails to drive premature cell cycle entry or premature transcription of *CLN1* (Nasmyth and Dirick, 1991; Wijnen *et al*., 2002). However, we found that *swi6*Δ cells are barely viable and rapidly accumulate suppressor mutations. Furthermore, since Swi6 is a component of both SBF and MBF it controls the transcription of hundreds of genes. Amongst these genes are 16 transcription factors, including several that strongly influence cell cycle-dependent transcription later in the cell cycle (Horak *et al*., 2002). Transcription factors downstream of SBF and MBF also control the transcription of numerous genes involved in ubiquitin-dependent proteolysis, which plays essential roles in controlling the levels of cyclins and other proteins (Horak *et al*., 2002). Thus, loss of Swi6 causes pervasive and complex cascading effects upon the expression of thousands of genes, including many that play central roles in cell cycle control and protein turnover (Horak *et al*., 2002). Together, these considerations make it difficult to draw clear and rigorous conclusions based on the effects of *swi6*Δ.

No previous studies have investigated the effects of Cln3 on production of late G1 cyclin proteins in *swi6*Δ cells. Here, we tested whether loss of *SWI6* influences accumulation of Cln2 in *MET25pr-3XHA-CLN2* cells. In the presence of *SWI6*, *GAL1-CLN3* advanced expression of Cln2 protein, as expected (Figure 3). Loss of Swi6 caused a loss of periodic expression of Cln2 protein in *MET25pr-3XHA-CLN2* cells, and overexpression of *CLN3* had no effect on expression of Cln2 protein in *MET25pr-3XHA-CLN2 swi6*Δ cells. Together, these observations suggest that Swi6 influences post-transcriptional mechanisms that are required for periodic expression of Cln2 protein. The data further suggest that Cln3 modulates levels of Cln2 protein at least partly via mechanisms that are dependent upon the many genes whose transcription is controlled, directly or indirectly, by SBF. However, the complex and pleiotropic effects of *swi6*Δ on expression of thousands of genes make it difficult to draw clear conclusions from the data.

**Figure 3:**
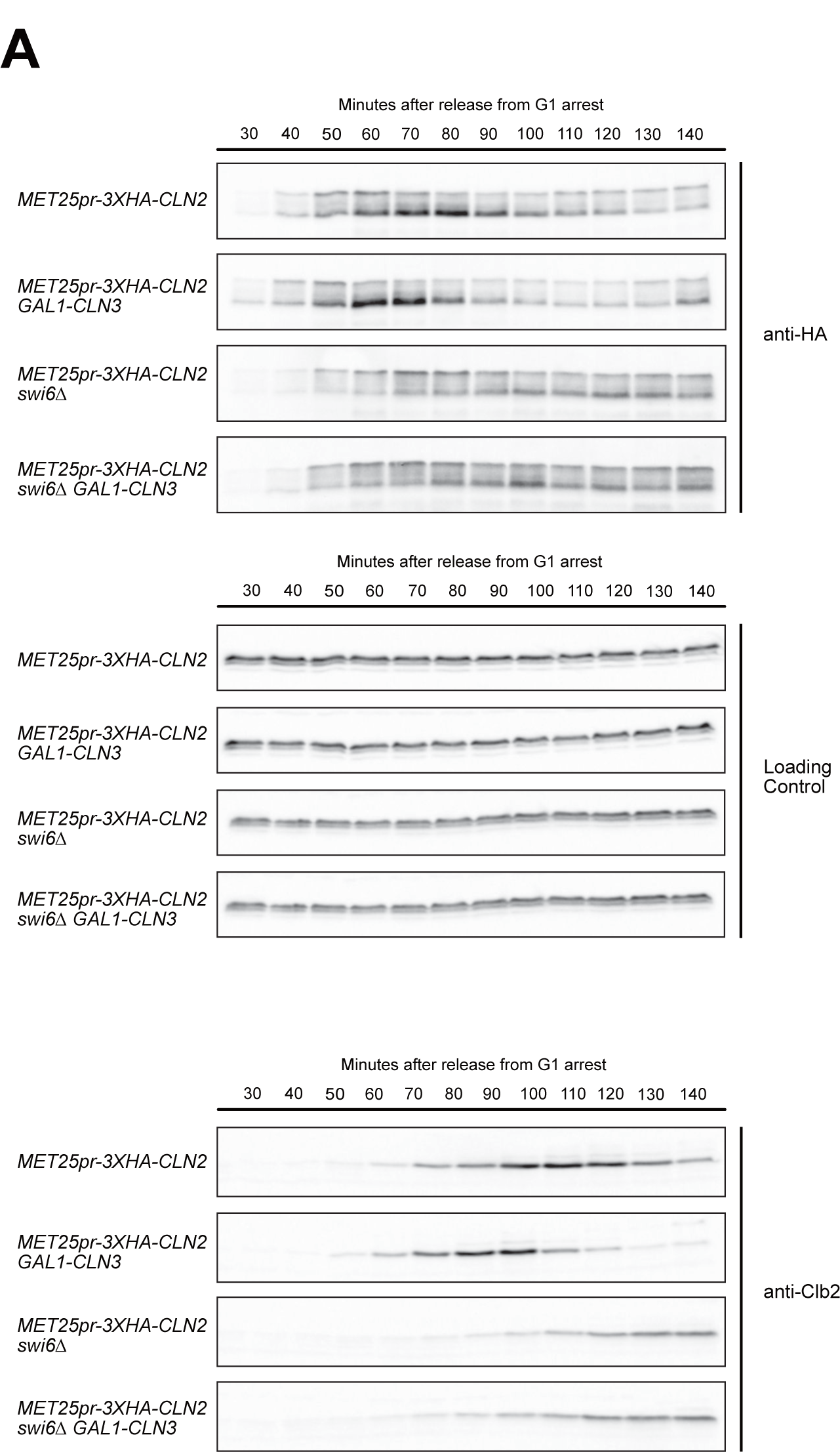
Cln3 requires Swi6 to influence Cln2 protein levels. **(A,B)** Cells were grown in YPGE medium overnight and synchronized in G1 phase using alpha factor. 2% galactose was added to the cultures 40 minutes prior to release from G1 arrest. Cells were released from G1 arrest into YP medium containing 2% galactose and samples were collected every 10 min. Western blotting was used to assay protein levels for Cln2 using 12CA5 mouse monoclonal anti-HA **(A)** and Clb2 using an anti-Clb2 antibody **(B)**.

Expression of Cln2 protein appeared to be repressed in early G1 phase in all four *MET25pr-3XHA-CLN2* strains, which suggests that post-transcriptional mechanisms repress accumulation of Cln2 protein in early G1 phase.

### Cln3 can influence cell size via mechanisms that are independent of transcriptional control of Cln1 and Cln2

Cln3 has been thought to influence cell size by regulating transcription of the late G1 cyclins (Cross, 1990; Tyers *et al*., 1993; Wijnen *et al*., 2002; Ferrezuelo *et al*., 2010). To further investigate, we tested whether overexpression of *CLN3* influences cell size in a strain in which normal transcriptional control of both *CLN1* and *CLN2* has been eliminated. To do this, we created a strain in which the *CLN1* gene was deleted and *CLN2* was controlled by the *MET25* promoter (*MET25pr-3XHA-CLN2 cln1*Δ). We then integrated a copy of the wild type *CLN3* gene under the control of the *GAL1* promoter. When grown in dextrose to repress expression of *GAL1-CLN3*, the *MET25pr-3XHA-CLN2 cln1*Δ *GAL1-CLN3* cells were larger than wild type (Figure 4A). When grown in 2% galactose, overexpression of *CLN3* drove a large decrease in the size of *MET25pr-3XHA-CLN2 cln1*Δ cells (Figure 4B). The fact that Cln3 can drive a decrease in cell size in *MET25pr-3XHA-CLN2 cln1*Δ cells shows that Cln3 can influence cell size via mechanisms that are independent of transcriptional control of *CLN1* and *CLN2*.

**Figure 4:**
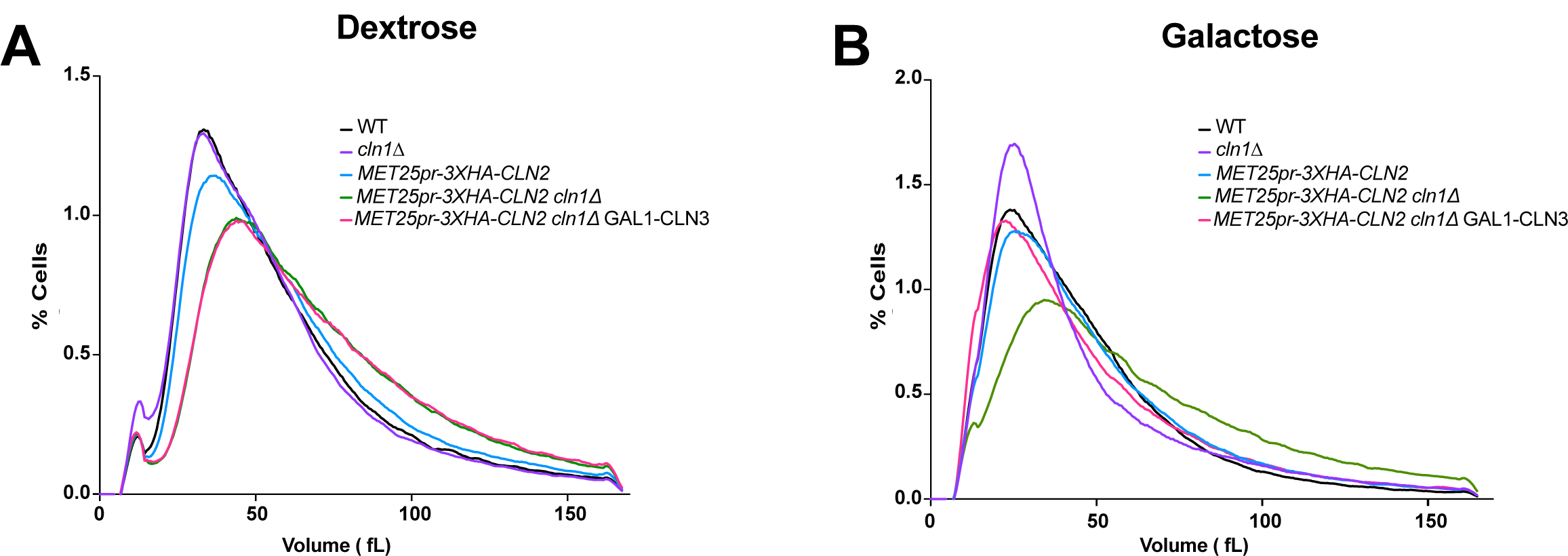
Cln3 can influence cell size via mechanisms that are independent of transcriptional control of Cln1 and Cln2. **(A, B)** Cells were grown in YPD medium or YP medium containing 2% galactose overnight to an OD_600_ between 0.4 - 0.6 and cell size was measured using a Beckman Coulter Counter Z2. Data shown represents the average of 3 biological replicates, where each biological replicate is the average of 3 technical replicates.

### Cln3 influences protein levels of targets of the SCF^Grr1^ ubiquitin ligase complex

The preceding results indicate that Cln3 can influence expression of Cln2 protein via post-transcriptional mechanisms. The only post-transcriptional mechanism currently known to play a major role in controlling Cln2 protein levels is ubiquitin-dependent proteolysis. Cln2 is targeted for ubiquitylation and proteolytic turnover by the SCF^Grr1^ ubiquitin ligase complex. Loss of SCF^Grr1^ activity causes a failure in Cln2 protein turnover, leading to accumulation of abnormally high levels of Cln2 and a loss of periodic expression of Cln2 protein (Willems *et al*., 1996; Skowyra *et al*., 1997). Thus, one potential explanation for the effects of Cln3 is that it influences activity of the SCF^Grr1^ complex. Consistent with this, previous studies have found that Cln3 can bind the SCF^Grr1^ complex. Moreover, *cln3*-Δ*177* binds less effectively to components of the SCF^Grr1^ complex compared to full length Cln3 (Willems *et al*., 1996; Landry *et al*., 2012), and we found that *cln3*-Δ*177* also appeared to be less effective than full-length *CLN3* at causing increased levels of Cln2 protein, consistent with a model in which Cln3 binds and inhibits SCF^Grr1^ (compare Figures 1B and 1D). To begin to test this model, we determined whether levels of other known SCF^Grr1^ targets are influenced by loss or gain-of-function of *CLN3*, which could support a model in which Cln3 influences the activity of the SCF^Grr1^ complex. Two of the best characterized targets of SCF^Grr1^ are Hof1 and Ndd1. Hof1 controls cytokinesis, whereas Ndd1 is an essential transcription factor that controls expression of a cluster of mitotic genes that includes mitotic cyclins (Blondel *et al*., 2005; Li *et al*., 2006; Edenberg *et al*., 2015). Both show strong cell cycle-dependent changes in protein levels.

We first analyzed Hof1-3XHA levels in asynchronous rapidly growing cells. Levels of Hof1 were decreased in *cln3*Δ cells and increased in *cln3*-Δ*177* cells (Figure 5A). In synchronized cells, *cln3*-Δ*177* led to an increase in Hof1 levels during mitosis, whereas *cln3*Δ led to a decrease in Hof1 levels (Figure 5B). We next compared the timing of expression of Hof1-3XHA and Cln3-6XHA in synchronized cells. Since the two proteins migrate at different locations in SDS-PAGE we were able to analyze levels of Hof1-3XHA and Cln3-6XHA in the same western blot. We also analyzed levels of the mitotic cyclin Clb2 as a marker for mitotic progression. Hof1-3XHA began to accumulate in early mitosis and reached peak levels late in mitosis, as levels of Clb2 began to decline (Figure 5C). Hof1 protein levels were strongly correlated with Cln3 protein levels, consistent with the possibility that Cln3 modulates levels of Hof1. The 6XHA tag used to detect Cln3 shows much higher sensitivity than the 3XHA tag used to detect Hof1, so relative levels of the two proteins cannot be compared.

**Figure 5:**
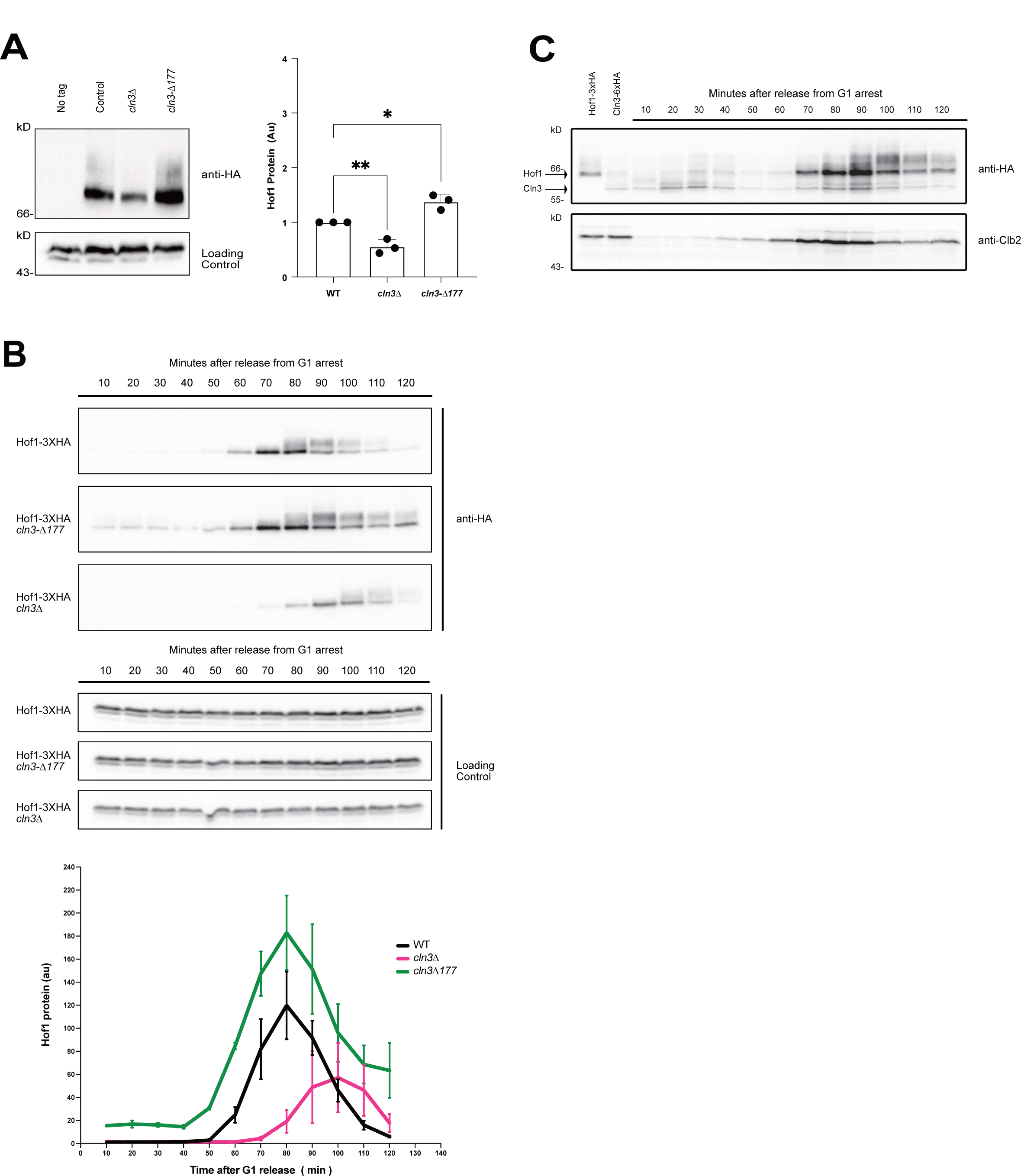
Cln3 influences levels of Hof1 protein. **(A)** Western blots showing levels of Hof1 protein in asynchronous wt, *cln3*Δ, and *cln3*-Δ*177* cells. Bar graphs show quantification of three biological replicates. Error bars show standard error of the mean. **(B)** Cells were grown overnight in YPD medium and synchronized in G1 phase using alpha factor. Cells were released from G1 arrest and samples were collected every 10 minutes. The behavior of Hof1-3XHA was assayed by western blot using 12CA5 mouse monoclonal anti-HA. An anti-Nap1 antibody served as a loading control. The graph shows quantification of data from three biological replicates normalized to loading control. Error bars represent the standard error of the mean. **(C)** Wild type cells containing Cln3-6XHA and Hof1-3XHA were grown overnight in YPD medium and sychronized in G1 phase with alpha factor. After release from the G1 arrest, the behavior of Cln3-6XHA and Hof1-3XHA was assayed by western blot using 12CA5 mouse monoclonal anti-HA.

We carried out a similar analysis for Ndd1. As with Hof1, *cln3*Δ caused a decrease in Ndd1 protein levels in both asynchronous and synchronous cells, whereas *cln3*-Δ*177* caused an increase in Ndd1 levels (Figures 6A,B). In contrast to Hof1, Ndd1 accumulates slightly before mitosis, consistent with its essential role in induction of transcription of mitotic cyclins and other mitotic genes. Therefore, Ndd1 levels are not closely correlated with Cln3 levels, and the effects of Cln3 on Ndd1 protein levels could be an indirect consequence of increased levels of late G1 cyclins, which are known to initiate mitotic transcription programs.

**Figure 6:**
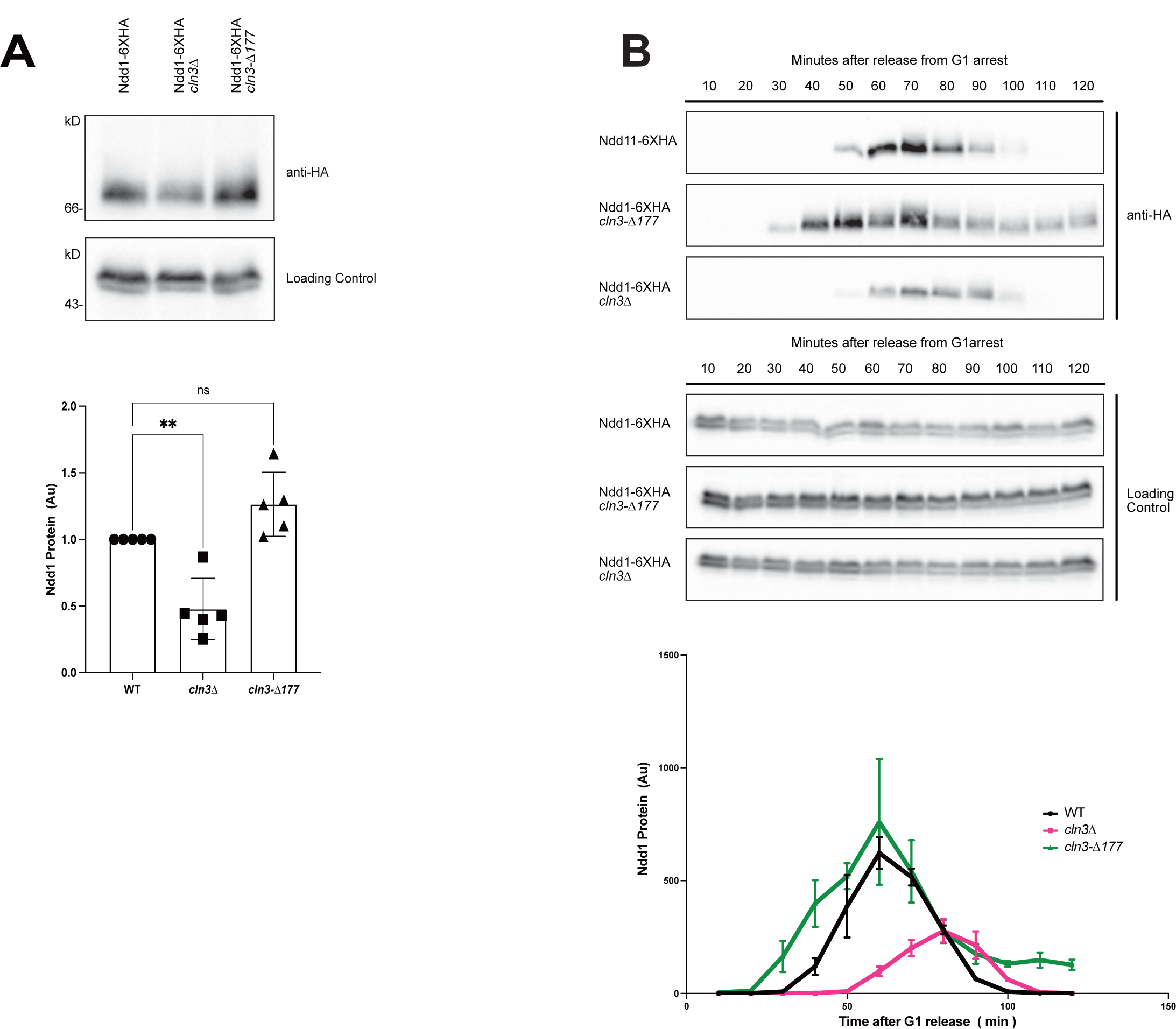
Cln3 influences levels of Ndd1 protein. **(A)** Western blots showing levels of Ndd1 protein in asynchronous wt, *cln3*Δ, and *cln3*-Δ*177* cells using 12CA5 mouse monoclonal anti-HA. Bar graphs show quantification of three biological replicates normalized to loading control. Error bars show standard error of the mean. **(B)** Cells were grown overnight in YPD medium and synchronized in G1 phase using alpha factor. Cells were released from G1 arrest and samples were collected every 10 minutes. The behavior of Ndd1-3XHA was assayed by western blot. An anti-Nap1 antibody served as a loading control. The graph shows quantification of data from three biological replicates. Error bars represent the standard error of the mean.

At the least, these results demonstrate that loss or gain-of-function of Cln3 causes complex effects on the proteome that make it difficult to draw simple conclusions about the effects of Cln3 mutants. Moreover, these results are consistent with the idea that Cln3 influences the activity of SCF^Grr1^, but do not rule out alternative models. Extensive additional work will be needed to further test whether Cln3 can influence the activity of the SCF^Grr1^ complex.

### Cln3 carries out functions in mitosis that influence cell size

Cln3 has been thought to exert all its effects in G1 phase. However, there is a second peak of Cln3 protein in mitosis (Landry *et al*., 2012; Zapata *et al*., 2014; Litsios *et al*., 2019), and we found that Cln3 can influence the expression of the Hof1 and Ndd1 proteins, which function after G1 phase. Together, these observations suggest that Cln3 could have functions when it appears later in the cell cycle during mitosis. Since most growth of budding yeast cells occurs during mitosis (Leitao and Kellogg, 2017), a mitotic function of Cln3 could make a major contribution to the effects of Cln3 on cell size. To investigate further, we used microscopy to analyze how loss- or gain-of-function of Cln3 influences the duration and extent of bud growth in mitosis. Since effects of Cln3 on cell size in G1 phase could influence growth in mitosis, we used conditional alleles to inactivate or overexpress Cln3 after cells passed through G1 phase. Conditional overexpression of Cln3 was achieved by expression of *CLN3* from the inducible *GAL1* promoter. Conditional inactivation of Cln3 was achieved with an auxin inducible degron version of *CLN3* (*cln3-AID*) (Nishimura *et al*., 2009). The *cln3-AID* allele caused a modest increase in cell size in the absence of auxin, which indicated that the AID tag caused decreased function of Cln3. Prolonged growth of *cln3-AID* cells in the presence of auxin caused a larger increase in cell size, although the increase was not as large as the increase caused by *cln3*Δ, which indicated that the *cln3-AID* allele caused a partial loss of function of Cln3 (Figure 7 – Figure Supplement 7).

*GAL1-CLN3* cells were arrested in G1 phase in media containing 2% glycerol and 2% ethanol to repress the *GAL1* promoter. Cells were released from the arrest and galactose was added to induce expression of *CLN3* when 15% of the cells had undergone bud emergence. To analyze loss of function, *cln3-AID* cells were released from a G1 arrest and auxin was added when 40-50% of the cells had undergone bud emergence. We induced gain- or loss-of-function of *CLN3* at different times relative to bud emergence because auxin-induced destruction of AID-tagged proteins occurs relatively rapidly (10-15 minutes), whereas high-level expression of proteins from the *GAL1* promoter occurs somewhat more slowly (20-30 minutes).

The durations of metaphase and anaphase were analyzed in single cells using a fluorescently tagged spindle pole protein, as previously described (Leitao and Kellogg, 2017; Jasani *et al*., 2020). The spindle poles in wild type control cells were tagged with mCherry, while the spindle poles in *cln3-AID* and *GAL1-CLN3* were tagged with GFP, which allowed analysis of control and experimental cells simultaneously under identical conditions. Cell growth was analyzed by measuring the size of the daughter bud as a function of time.

Overexpression of Cln3 caused a decrease in the durations of both metaphase and anaphase, as well as a decrease in the size at which daughter buds complete metaphase and anaphase (Figures 7A,B). Conversely, loss of function of Cln3 caused an increase in the duration of metaphase and an increase in daughter bud size at the end of metaphase (Figures 7C,D). Loss of function of Cln3 did not cause a statistically significant change in the duration of anaphase.

**Figure 7:**
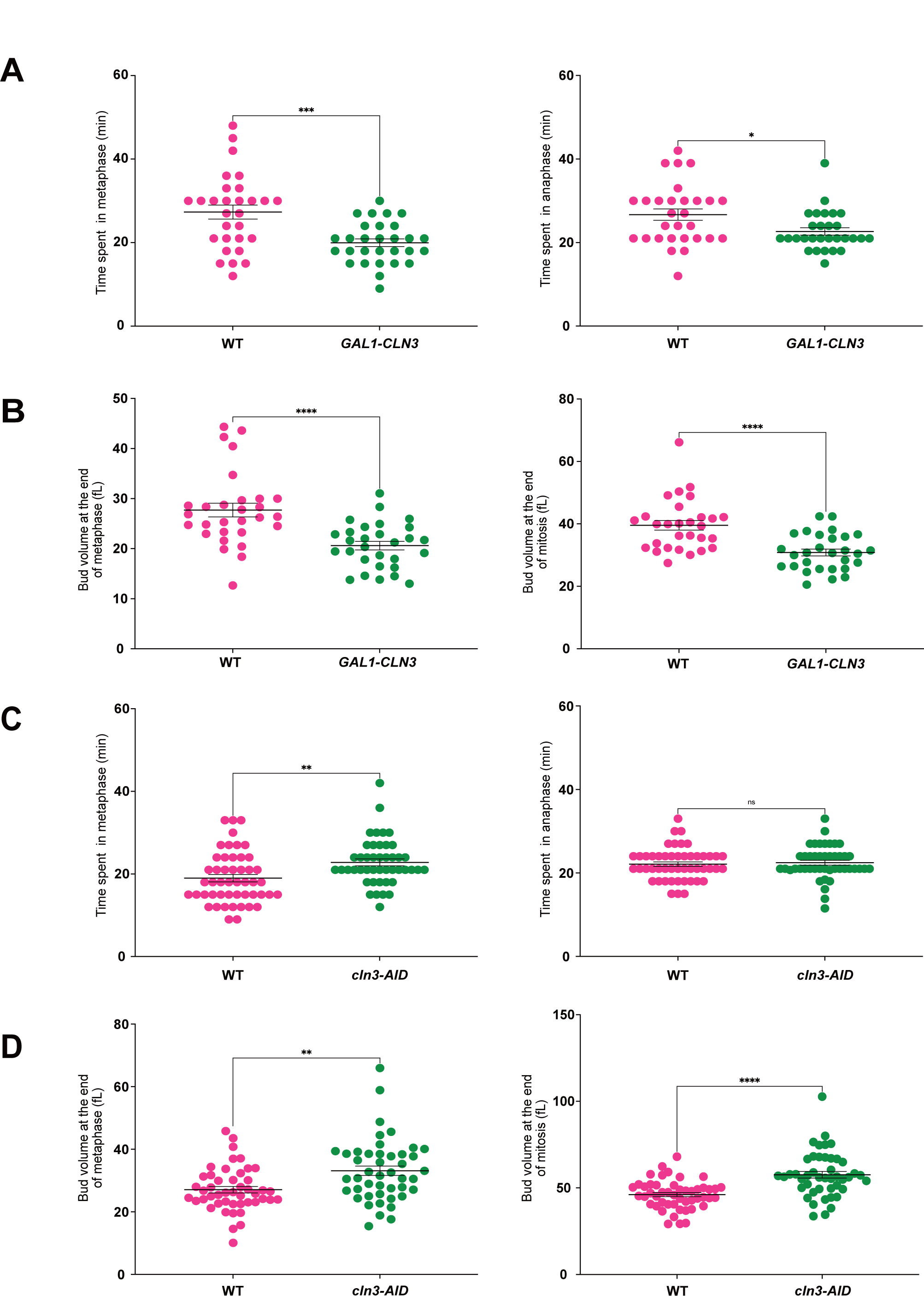
Cln3 carries out functions in mitosis that influence cell size. Mitotic spindle poles in wild type control cells were labelled with Spc42-mRUBY2, whereas the mitotic spindle poles of *GAL1-CLN3* or *cln3-AID* cells were labeled with Spc42-eGFP. For experiments comparing wild type to *GAL1-CLN3*, cells were grown in complete synthetic media (CSM) containing 2% glycerol and 2% ethanol overnight. The cells were then arrested in G1 phase using alpha factor and wild type and mutant cells were mixed before releasing from the arrest. Cells were released in CSM containing 2% glycerol/ethanol and 2% galactose was added 60 minutes after release. For experiments comparing wild type cells to *cln3-AID*, cells were grown overnight in CSM containing 2% dextrose and were arrested in G1 phase using alpha factor. The wild type and mutant cells were mixed before releasing from the G1 phase arrest. 0.5 mM auxin was added at 50 minutes after release. Cells were imaged by confocal microscopy at 3-minute intervals at a constant temperature of 27°C. **(A)** Graphs comparing the duration of each stage of mitosis in wild type and *GAL1-CLN3* cells. **(B)** Graphs comparing bud volume at the end of metaphase and the end of mitosis in wild type and *GAL1-CLN3* cells. **(C)** Graphs comparing the duration of each stage of mitosis in wild type and *cln3-AID* cells. **(D)** Graphs comparing bud volume at the end of metaphase and the end of mitosis in wild type and *cln3-AID* cells. Bars indicate the mean and the standard error of the mean for n=29 (wild type, *GAL1CLN3*) in panels A and B, and n=49 (wild type) and n=45 (*cln3-AID)* in panels C and D.

Together, these data show that Cln3 can influence the duration and extent of growth in mitosis, which suggests Cln3 has functions outside of G1 phase that strongly influence cell size. The data are consistent with our finding that Cln3 can influence expression of proteins outside of G1 phase. Previous studies have shown that loss of Hof1 causes a large increase in cell size, which suggests that decreased expression of Hof1 could contribute to the large size of *cln3*Δ cells. However, the mechanisms by which Cln3 influences cell growth and size in mitosis remain largely unknown.

## Discussion

### Expression of late G1 phase cyclin proteins is strongly influenced by post-transcriptional mechanisms

Current models for cell cycle entry suggest that transcriptional mechanisms play a critical role in cell cycle-dependent expression of Cln2 protein, and that Cln3 influences cell cycle entry solely via transcriptional mechanisms that initiate expression of Cln2 and other late G1 phase cyclins (Tyers *et al*., 1993; Dirick *et al*., 1995; Costanzo *et al*., 2004; de Bruin *et al*., 2004; Kõivomägi *et al*., 2021). Here, we carried out new tests of these models. Importantly, we analyzed production of Cln2 protein as the critical output of mechanisms that drive cell cycle entry. We found that Cln3 and Whi5, which are thought to be critical regulators of *CLN2* transcription and cell cycle entry, are not required for cell cycle-dependent expression of Cln2 protein. We also found that Cln2 protein expression shows strong cell cycle-dependent periodicity even when it is expressed from heterologous promoters that are not regulated by SBF, which indicates that post-transcriptional mechanisms play an important role in enforcing cell cycle-dependent expression of Cln2. Finally, we found that Cln3 can influence production of Cln2 protein when Cln2 is expressed from heterologous promoters, which indicates that Cln3 can influence Cln2 protein levels via post-transcriptional mechanisms. Together, these surprising discoveries show that post-transcriptional mechanisms play a major role in regulation of Cln2 expression.

Two independent studies detected Cln3 at SBF promoters via chromatin immunoprecipitation assays, which suggests that Cln3 carries out functions at SBF promoters (Wang *et al*., 2009; Kõivomägi *et al*., 2021). A recent study concluded that Cln3/Cdk1 functions at SBF promoters to directly phosphorylate and activate RNA polymerase (Kõivomägi *et al*., 2021). This model was supported by previous studies in which it was found that loss of *CLN3* in an otherwise wildtype background causes reduced transcription of *CLN2* (Cross and Tinkelenberg, 1991; Tyers *et al*., 1993; Di Como *et al*., 1995; Dirick *et al*., 1995). Here, we found that loss of *CLN3* does not cause a reduction in peak Cln2 protein levels in *whi5*Δ cells, which is inconsistent with the idea that Cln3 acts independently of Whi5 to promote activation of RNA polymerase at the *CLN2* promoter. We also found that cells that lack both Whi5 and Cln3 express <u>more</u> Cln2 protein than wild type cells, and the timing of Cln2 expression is indistinguishable from wild type cells (Figure 1B). Together, the data are more consistent with previous genetic data that led to a model in which the primary function of Cln3 at the *CLN2* promoter is to drive release of Whi5 from SBF (Costanzo *et al*., 2004; de Bruin *et al*., 2004). The available data do not rule out the possibility that Cln3 drives release of Whi5 via mechanisms that work at the post-transcriptional level to promote production of more Cln2/Cdk1, which is thought to directly phosphorylate proteins at the Cln2 promoter to drive release of Whi5. Loss of Cln3 caused a slight delay in Cln2 protein expression in *whi5*Δ cells compared to *whi5*Δ alone, which could be due entirely to post-transcriptional mechanisms (Figure 1B). Overall, however, the mechanisms by which Cln3 drives release of Whi5 remain poorly understood.

The key experiments that led to a model in which Cln3/Cdk1 phosphorylates RNA polymerase were based on experiments that utilized engineered Cln3-Cdk1 fusion proteins (Kõivomägi *et al*., 2021). The fusion protein approach was used because it has not been possible to purify a Cln3/Cdk1 complex, which is surprising because other cyclins bind Cdk1 to form a tight stoichiometric complex that can only be dissociated under denaturing conditions. The fact that it has not been possible to purify a Cln3/Cdk1 fusion protein indicates that Cln3 binds Cdk1 with dramatically lower affinity. Consistent with this, one well-controlled study did not detect any Cdk1 activity associated with Cln3 (Schneider *et al*., 2004), whereas several others detected only very weak activity (Tyers *et al*., 1992, 1993; Levine *et al*., 1996; Miller *et al*., 2005). The fact that Cln3 binds Cdk1 with dramatically lower affinity raises the possibility that the Cdk1 in a Cln3-Cdk1 fusion protein could form complexes with other cyclins, which bind Cdk1 with a much higher affinity. Thus, Cln3-Cdk1 fusion proteins could form highly active Cln3-Cdk1/cyclin complexes and recruit them to the *CLN2* promoter via the known interaction of Cln3 with SBF promoters. In this case, the recruitment of ectopic highly active Cdk1/cyclin complexes to the *CLN2* promoter would be expected to robustly phosphorylate and activate RNA polymerase because it is well known that there are Cdk/cyclin complexes that have evolved to specifically control transcription via phosphorylation of the RNA polymerase (Lim and Kaldis, 2013). Similarly, the activity of Cln3-Cdk1 fusion proteins in vitro could represent the sum activity of multiple diverse Cln3-Cdk1/cyclin complexes. Additional controls will be necessary to rule out these kinds of scenarios and to more clearly establish the role of Cln3 at SBF promoters.

The discovery that Cln3 can influence production of Cln2 protein via post-transcriptional mechanisms has important implications, as it expands the range of possible models for control of cell cycle entry and cell size. For example, previous studies have shown that Cln2 promotes its own transcription via a positive feedback loop in which Cln2/Cdk1 is thought to directly phosphorylate and regulate proteins at the *CLN2* promoter (Nasmyth and Dirick, 1991; Costanzo *et al*., 2004; de Bruin *et al*., 2004; Skotheim *et al*., 2008; Wagner *et al*., 2009). Thus, an ability of Cln3 to directly influence Cln2 protein levels could determine when the positive feedback loop is engaged to drive full entry into the cell cycle. Since Cln3 accumulates gradually during G1 phase in a manner that is dependent upon and proportional to cell growth and nutrient availability (Litsios *et al*., 2019; Sommer *et al*., 2021), Cln3 could influence the amount of growth required to trigger cell cycle entry, which would provide a mechanism for linking cell cycle entry to cell growth and nutrient availability.

A number of models could explain how post-transcriptional mechanisms help enforce periodic expression of late G1 phase cyclins. The rise in Cln2 levels in G1 phase could be explained by mechanisms that inhibit SCF^Grr1^. For example, signals related to cell growth could inhibit SCF^Grr1^ to drive an increase in Cln2 protein levels. If Cln3 were to influence the activity of SCF^Grr1^, it could influence the amount of growth needed to initiate expression of Cln2 protein. The decline in Cln2 levels at the end of G1 phase could be explained by a model in which cyclin/Cdk complexes expressed after cell cycle entry relay signals that inhibit the SCF^Grr1^ complex.

Numerous models are possible at this point and considerable additional work will be required to gain a better understanding of the mechanisms that drive cell cycle-dependent expression of Cln2 protein and cell cycle entry. The mechanisms that drive cell cycle entry in mammalian cells also remain poorly understood (Rubin *et al*., 2020). Thus, a better understanding of cell cycle entry in yeast could lead to the discovery of conserved mechanisms that are relevant to mammalian cells.

### Cln3 is likely to have pervasive effects on the proteome

The only major post-transcriptional mechanism that is known to regulate Cln2 protein levels is ubiquitin-dependent proteolysis, which is mediated by the SCF^Grr1^ ubiquitin ligase complex. Therefore, the most straightforward hypothesis for how Cln3 influences Cln2 levels is via regulation of SCF^Grr1^. To begin to test this idea, we determined whether Cln3 can influence the levels of other known protein targets of SCF^Grr1^. We found that protein levels of two of the best characterized targets of SCF^Grr1^ (Hof1 and Ndd1) are strongly influenced by Cln3 in a manner similar to Cln2. These observations are consistent with the idea that Cln3 influences Cln2 protein levels via SCF^Grr1^, but do not rule out alternative models. At the least, the data show that Cln3 is likely to have pervasive effects on the proteome, which suggests that the effects of Cln3 on cell size could be due to complex effects on the expression of multiple proteins. The fact that Cln3 modulates levels of multiple SCF^Grr1^ targets could explain the genetic data that indicate that Cln3 can influence cell size via targets other than Whi5 (Costanzo *et al*., 2004; Wang *et al*., 2009).

### Evidence for mitotic functions of Cln3 that influence cell size

Influential studies carried out over 50 years ago reached the conclusion that cell size is regulated primarily in G1 phase in budding yeast (Hartwell and Unger, 1977). However, more recent work has shown that little growth occurs in G1 phase (Leitao and Kellogg, 2017). For example, cell volume increases by only 10-20% during G1 phase when cells are growing in rich nutrients. Rather, most growth takes place during growth of the daughter bud, which occurs almost entirely during mitosis (Leitao and Kellogg, 2017). Moreover, the extent of bud growth during mitosis is modulated by nutrient availability, which has large impacts on daughter cell size. There is evidence for nutrient-modulated mechanisms that measure the extent of bud growth to ensure that sufficient growth has occurred before cytokinesis (Leitao and Kellogg, 2017; Leitao *et al*., 2019; Jasani *et al*., 2020).

The idea that cell size is regulated primarily in G1 phase has strongly influenced current models for the functions of Cln3, which assume that Cln3 exerts its effects on cell size solely in G1 phase. However, the idea that Cln3 functions solely in G1 phase is challenged by several observations. First, there is a second peak of Cln3 protein in mitosis, which suggests that Cln3 has functions in mitosis (Landry *et al*., 2012; Zapata *et al*., 2014; Litsios *et al*., 2019). Second, analysis of Coulter Counter data shows that overexpression of Cln3 causes daughter cells to be born at a dramatically reduced size, which can only occur if daughter buds undergo less growth before mitosis (Costanzo *et al*., 2004; Zapata *et al*., 2014). These observations led us to hypothesize that Cln3 could influence cell size via effects on the duration and extent of bud growth in mitosis. To test this, we developed methods to induce conditional loss or gain-of-function of Cln3 after G1 phase and before mitosis, which showed that loss of Cln3 causes daughter cells to be born at a larger size, while overexpression of Cln3 causes them to be born at a smaller size. These data indicate that Cln3 can influence the duration and extent of bud growth in mitosis, and that mitotic functions of Cln3 can not be ignored when considering models for how Cln3 influences cell size.

The targets of Cln3 in mitosis that influence cell size are unknown. We found that Cln3 influences levels of Hof1, a mitotic regulator of cytokinesis, and that Hof1 protein levels in mitosis are correlated with Cln3 protein levels, consistent with the idea that Cln3 has functions in mitosis. We and others have found that *hof1*Δ cells are larger than wild type cells (Li *et al*., 2006) and we report here that loss of Cln3 leads to a reduction in Hof1 protein levels. Together, these observations suggest that part of the effects of Cln3 on cell size could be due to effects on Hof1 protein levels; however, Hof1 has not previously been implicated in cell size control and we found that overexpression of HOF1 does not reduce cell size. Additional work will be needed to gain a better understanding of how Cln3 influences cell size in mitosis.

## Materials and Methods

### Yeast strains, plasmids, media and cell cycle time courses

All strains are in the W303 background and carry a deletion of the BAR1 gene to facilitate arrest with alpha factor (MATa *leu2-3,112 ura3-1 can1-100 ade2-1 his3-11,15 trp1-1 GAL+, ssd1-d2 bar1-*). The additional genetic features of the strains are listed in Table 3. Genetic alterations, such as epitope tagging, promoter swaps, and gene deletions were carried out using homologous recombination at the endogenous locus (Longtine *et al*., 1998; Janke *et al*., 2004).

Plasmid pAB1 was used to integrate *GAL1-CLN3-3XHA* at the *URA3* locus. To create pAB1, the *GAL1* promoter was amplified and cloned into the KpnI and Xho1 sites of pRS306 (primers: CGCGGTACCTTATATTGAATTTTCAAAAATTCT and GCGCCTCGAGTATAGTTTTT-TTCTCCTTGACG) to make pDK20. A 3XHA tag sequence was then cloned into the EagI and SacII sites of pDK20 to create pSH32. The *CLN3* coding sequence was amplified and cloned into the XhoI and EagI sites of pSH32 (primers: GCGCTCGAGATGGCCATATTGAAGGATAC-CATAATTAGATACGC and CGCCGGCC-GGCGAGTTTTCTTGAGGTTGCTACTATC). pAB1 is cut with Stu1 to target integration at the URA3 locus.

To create a plasmid that allows PCR-based replacement of promoters with the *YPK1* promoter, the *YPK1* promoter was amplified by PCR with BglII and PacI sites and was used to replace the *GAL1* promoter in plasmid pFA6a-His3MX6-pGAL with the *YPK1* promoter (pDK132, primers: GCGAGATCTGATGTTTTAACTGATCTTAATTTATATGTAGAGGA, GCGTTAATTAATTTCAGGAACTGTATTAATGTTTGTTGATAT).

For cell cycle time courses, cells were synchronized with 0.5 ug/ml alpha factor between 3-3.5 hours at room temperature and released from the arrest by washing cells 3 times with 25 mls of growth medium. For cell cycle time courses where *GAL1-CLN3* was induced, cells were grown in YP medium containing 2% glycerol and 2% ethanol (YPGE) and were arrested with alpha factor for 3.5-4 hours at room temperature. In Figure 7, *GAL1-CLN3* was induced by addition of 2% galactose at 60 minutes after release from alpha factor arrest. In Figure 1C, 2C and Figure 3, 2% galactose was added to the cultures 40 minutes before releasing from the arrest and cells were then released from the arrest into YP medium containing 2% galactose. All time courses were done at 30°C.

### Cell size analysis

Cell size was measured with a Beckman Coulter Channelyzer Z2. Briefly, cells were grown in 10 mls culture medium to an OD_600_ between 0.4 - 0.6. Cells were then fixed by addition of 1/10 volume of formaldehyde for 30-60 minutes. Cells were pelleted and resuspended in 500 µl of PBS, 0.02% sodium azide and 0.1% Tween-20 and briefly sonicated. Cell size was measured using a Coulter Channelizer Z2; Beckman Coulter). In each Figure, cell size data represents the average of 3 biological replicates, in which each biological replicate is the average of 3 technical replicates.

### Single cell analysis of cell growth and size during the cell cycle

Analysis of cell growth and size during the cell cycle was carried out as previously described (Leitao and Kellogg, 2017; Jasani *et al*., 2020). Briefly, cells were grown in complete synthetic media (CSM) containing 2% dextrose (CSM-Dex) or 2% glycerol and 2% ethanol (CSM-G/E). For conditional expression of *GAL1-CLN3* 2% Galactose was added to the media when cells reached approximately 15% bud emergence, indicating that they were 20-30 minutes away from entering mitosis, which allowed sufficient time for expression *CLN3* from the *GAL1* promoter before mitosis. For conditional degradation of Cln3 in mitosis, auxin was added to the medium once cells were approximately 40-50% budded to allow for degradation of Cln3 before the beginning of mitosis. Sample preparation, data acquisition and processing was performed as previously described by (Leitao and Kellogg, 2017; Jasani *et al*., 2020).

### Western Blotting

Western Blotting was done as previously described (Sommer *et al*., 2021). All polyacrylamide gels were 10% polyacrylamide and 0.13% bis-acrylamide, except for gels used for Cdk1 protein analysis, which were 12.5% polyacrylamide with 0.10% bis-acrylamide. All gels were transferred to a nitrocellulose membrane using Bio-Rad Trans-Blot Turbo Transfer system. Blots were probed overnight at 4°C in 3% dry milk in western wash buffer (PBS + 250 mM NaCl +0.1% Tween-20) using 12CA5 mouse monoclonal anti-HA, anti-Nap1, anti-Clb2, anti-GST or anti-Cdk1. Primary antibodies were detected with an HRP-conjugated donkey anti-rabbit secondary antibody or HRP-conjugated donkey anti-mouse secondary antibody as previously described by incubating for 1 hour at room temperature. Western blots were imaged using ECL reagents (K-12045-D50; Advansta) and a Bio-Rad Chemi-Doc Imaging system.

### Western Blotting Quantification

All quantifications were done using Image Lab (Bio-Rad) as previously described (Jasani *et al*., 2020; Sommer *et al*., 2021). For alpha factor block and release experiments, the signal at each time point was calculated as a ratio over the signal at the 10-minute time point for the control strain. The signal for each time point was then normalized to the loading control. For log phase samples, signals were calculated as a ratio over the WT control and then normalized to loading control.

## Experimental Replicates

All experiments were repeated for a minimum of 3 biological replicates. Biological replicates are defined as experiments carried out on different days with cultures started with newly thawed cells.

## Materials Availability

All yeast strains, plasmids and antibodies described in this study are available upon request.

## Data Availability

All source data are included within the manuscript files.

## Acknowledgements

This work was supported by NIH grant GM131826. A.B. was supported by NIH grants GM058903 and GM133391, and by the ARCS Foundation, Marilyn D. Davis Memorial Scholarship, Denice Denton Award for Promoting Diversity, Café Bustelo El Café Del Futuro Scholarship, and UCSC Slug Support Office. We thank Benjamin Abrams (University of California, Santa Cruz) for providing technical support for the microscopy experiments, and David Toczyski (University of California, San Francisco) for providing 12CA5 mouse monoclonal anti-HA antibody.

---

**Figure 7, figure supplement:**
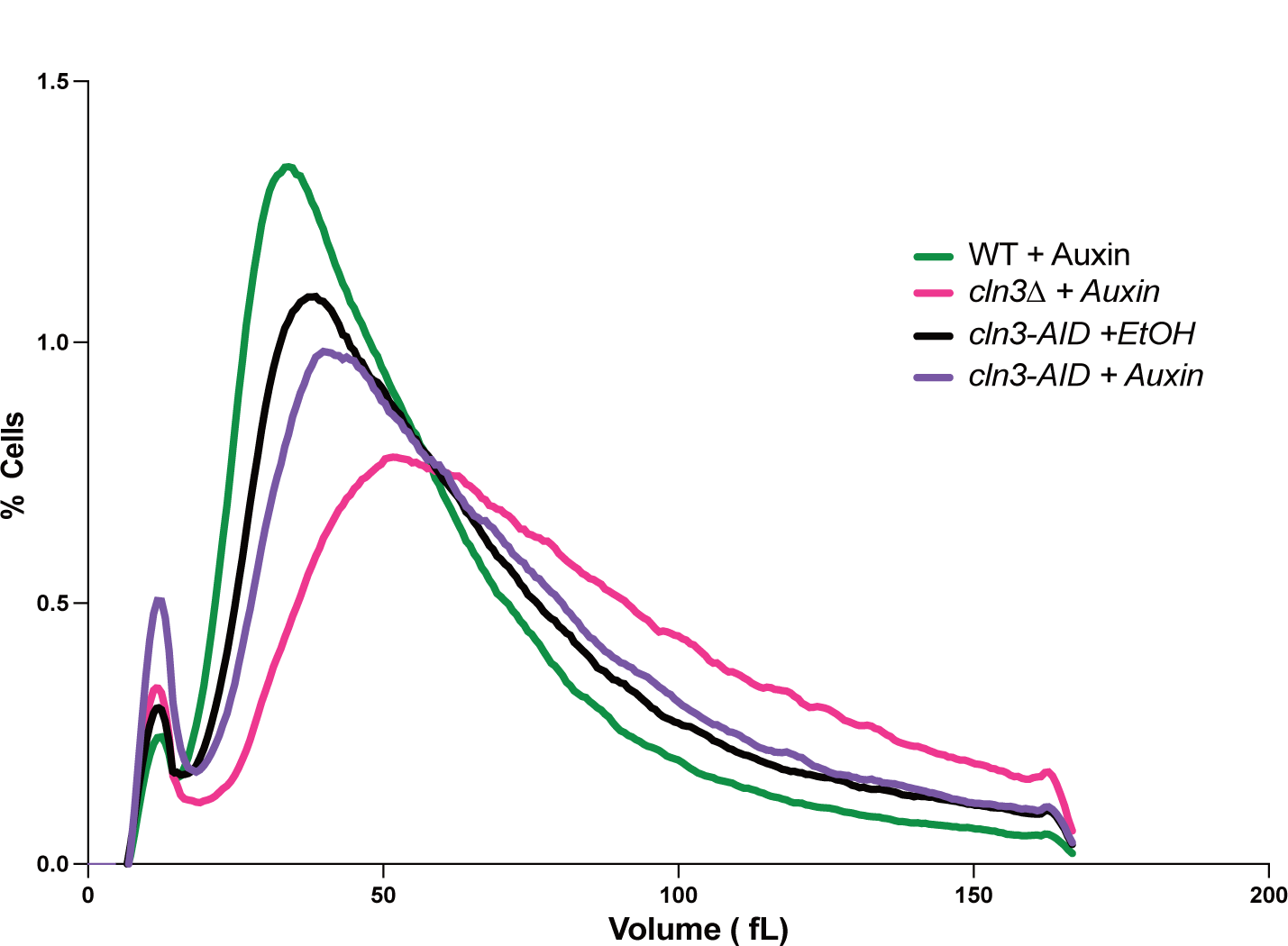
Cell size analysis of *cln3-AID* cells. Cells were grown overnight in YPD medium containing either 0.5 mM auxin or an ethanol solvent control. Cell size was measured using a Coulter counter. Data shown represents the average of 3 biological replicates, in which each biological replicate represents the average of 3 technical replicates.

**Table 1:** Strains used in this study.

| Strain | Genotype | Figure |
| --- | --- | --- |
| DK1295 | *whi5Δ::KanMx4* | Figure 1 |
| DK4074 | *CLN2-3XHA::hphNT1* | Figure 1 |
| DK4102 | *CLN2-3XHA::hphNT1 whi5Δ::KanMX4* | Figure 1 |
| DK4108 | *CLN2-3XHA::hphNT1 whi5Δ::KanMX4 cln3-Δ177-6XHA::HIS3MX6* | Figure 1 |
| DK4114 | *CLN2-3XHA::hphNT1 whi5Δ::KanMX4, cln3Δ::His3MX6* | Figure 1 |
| DK4125 | *CLN2-3XHA::hphNT1 whi5Δ::KanMX4, GAL1-CLN3::URA3* | Figure 1 |
| DK4032 | *MET25pr-3XHA-CLN2::NatNT2 cln3Δ::HIS3* | Figure 2 |
| DK4034 | *MET25pr-3XHA-CLN2::NatNT2 cln3-Δ177-6XHA::HIS3MX6* | Figure 2 |
| DK4026 | *MET25pr-3XHA-CLN2::NatNT2* | Figure 2, 3 |
| DK4714 | *His3-YPK1pr-CLN2-9XMyC::hphNT1* | Figure 2 |
| DK4718 | *His3-YPK1pr-CLN2-9xMyC::hphNT1 GAL1-CLN3::URA3* | Figure 2 |
| DK4527 | *MET25pr-3XHA-CLN2::NatNT2 GAL1-CLN3::URA3* | Figure 3 |
| DK4545 | *MET25pr-3XHA-CLN2::NatNT2 swi6Δ::HIS3* | Figure 3 |
| DK4548 | *MET25pr-3XHA-CLN2::NatNT2 swi6Δ::HIS3 GAL1-CLN3::URA3* | Figure 3 |
| DK4144 | *MET25pr-3XHA-CLN2::NatNT2 cln1Δ::hphNT1* | Figure 4 |
| DK4230 | *MET25pr-3XHA-CLN2::NatNT2 cln1Δ::hphNT1 GAL1-CLN3::URA3* | Figure 4 |
| KA65 | *cln1Δ::TRP* | Figure 4 |
| DK186 |  | Figure 4, Figure 7-supplement |
| DK3922 | *HOF1-3XHA::kITRP1* | Figure 5 |
| DK3945 | *HOF1-3XHA:: kITRP1 cln3-Δ177-6XHA::HIS3MX6* | Figure 5 |
| DK3960 | *HOF1-3XHA:: kITRP1 cln3Δ::His3MX6* | Figure 5 |
| DK3991 | *HOF1-3XHA:: kITRP1 Cln3-6XHA::His3MX6* | Figure 5 |
| DK4218 | *NDD1-6XHA::hphNT1* | Figure 6 |
| DK4233 | *NDD1-6XHA::hphNT1 cln3Δ::His3MX6* | Figure 6 |
| DK4235 | *NDD1-6XHA::hphNT1 cln3-Δ177-6XHA::His3MX6* | Figure 6 |
| DK3348 | *MYO1-GFP::TRP SPC42-GFP::His3MX6 GAL1-CLN3-3XHA::URA3* | Figure 7 |
| DK3510 | *his3::HIS3+TIR1 leu2::LEU2+TIR1 SPC42-mRuby2::KanMX4* | Figure 7 |
| DK3580 | *SPC42-yomRuby2::KanMX4 GAL1::URA3* | Figure 7 |
| DK3524 | *his3::HIS3+TIR1 leu2::LEU2+TIR1 cln3-AID::KanMX4 SPC42-GFP::hphNT1* | Figure 7, Figure 7-supplement |
| SH184 | *cln3Δ::His3MX6* | Supplemental Figure 7 |

